# Dynamic functional connectivity is modulated by the amount of p-Tau231 in blood in cognitively intact participants

**DOI:** 10.1101/2024.05.29.596323

**Authors:** Martín Carrasco-Gómez, Alejandra García-Colomo, Alberto Nebreda, Ricardo Bruña, Andrés Santos, Fernando Maestú

## Abstract

**INTRODUCTION:** Electrophysiology and plasma biomarkers are early and non-invasive candidates for Alzheimer’s disease detection. The purpose of this paper is to evaluate changes in dynamic functional connectivity measured with magnetoencephalography, associated with the plasma pathology marker p-tau231 in unimpaired adults.

**METHODS:** 73 individuals were included. Static and dynamic functional connectivity were calculated using leakage corrected amplitude envelope correlation. Each source’s strength entropy across trials was calculated. A data-driven statistical analysis was performed to find the association between functional connectivity and plasma p-tau231 levels. Regression models were used to assess the influence of other variables over the clusters’ connectivity.

**RESULTS:** Frontotemporal dynamic connectivity positively associated with p-tau231 levels. Linear regression models identified pathological, functional and structural factors that influence dynamic functional connectivity.

**DISCUSSION:** These results expand previous literature on dynamic functional connectivity in healthy individuals at risk of AD, highlighting its usefulness as an early, non-invasive and more sensitive biomarker.

## 1. Introduction

The biological conception of Alzheimer’s disease (AD) led to the definition of a continuum that begins with the appearance of the pathological hallmarks of the disease but no associated clinical symptom. These changes can emerge up to 20 years before the diagnosis, which highlights the importance of early detection tools to identify individuals in this state and track the progression of the disease, along with the effect of potential interventions [1].

Possitron emission tomography (PET) and cerebrospinal fluid (CSF) biomarkers are considered poor candidates to this end, due to their high cost, invasiveness, and low accessibility. Therefore, increasing attention has been placed on blood biomarkers [2]. In this vein, the recently discovered species of phosphorylated tau at threonine 231 (p-tau231) presents promising results. Abnormal levels, both in CSF and plasma, can be detected as early on as the preclinical stage. Furthermore, its levels parallel those of amyloid-beta (Aβ), tracking its changes even before Aβ-PET positivity has been achieved. Therefore, given the temporal and disease-specific association between both pathology markers, p-tau231 can be employed as a proxy of Aβ pathology [3–7].

Neuronal dysfunction and macroscopic functional changes can also be detected in this preclinical stage. In this context, electrophysiological measures are being examined as early biomarkers, given their minimal invasiveness and ability to detect incipient alterations in brain activity [8,9]. Studies addressing the changes in static functional connectivity (sFC) along the AD continuum reveal an inverted U-shaped pattern, whereby the preclinical stage is marked by increases in sFC, followed by a subsequent state of hypoconnectivity [10–14]. The initial state of hyperconnectivity has been associated with Aβ deposition, known to establish a positive feedback loop with neuronal hyperexcitability that underlies the observed hyperconnectivity [15–17]. Furthermore, recent studies have found interesting results when exploring sFC in the alpha subfrequency bands [14,18], with a special mention to García-Colomo et al. [10] where individuals at higher risk of AD showed increased sFC values, and the longitudinal increase across time in FC was associated with the Aβ and AD pathology proxy p-tau231.

Notably, resting-state activity has been broadly used in this context, in an attempt to study the organization of brain communication while avoiding inter-center bias and using comparable metrics between subjects with different levels of pathology [19]. Functional connectivity from resting-state recordings has been typically assessed by averaging the connectivity values across the whole recording, thus providing a value of sFC, which assumes that connections and their strength are stationary over time [20]. However, brain activity and networks are not static; rather, regions constantly connect and disconnect from each other in complex temporal dynamics even during resting state [21,22]. Therefore, what has been classically interpreted as inter-subject variability or background noise, might in fact be temporal variability. Thus, capturing the changes and fluctuations in connectivity beyond classical sFC can improve the understanding of brain function. This field, dedicated to the study of spatiotemporal dynamics of brain networks, is known as “dynamic functional connectivity” (dFC). Given the relevance of early biomarkers and the alterations described in sFC as early as the preclinical stage, studying and characterizing incipient changes in dFC can aid in the understanding of the development of AD and contribute to identify individuals entering the continuum.

Most studies involving dFC in the AD continuum have been conducted using functional magnetic resonance imaging (fMRI). These consistently report a progressive reduction of dFC among mild cognitive impairment and AD patients, involving relevant regions and networks such as the precunei and default mode network [23–25]. Studies addressing cognitively unimpaired individuals with AD pathology markers, however, show conflicting results, finding increases [26] and decreases [27] in dFC, this is, in the variability of FC over time, associated with the presence of Aβ. These clashing findings could be due to differences in methodology used to evaluate dFC and suggest an influence of AD pathology over dFC, which could be more sensitive to pathology from the earliest stages than sFC.

Although more appropriate to measure rapid dynamic changes in connectivity, electrophysiological studies are scarcer. Nonetheless, and similarly to fMRI, research performed in the dementia stage reveals a reduction of dFC, measured with different metrics such as variance and hidden Markov models, as well as related metrics such as entropy and dynamic changes in power [28,29]. Results are conflicting when considering mild cognitive impairment patients [22,30] and, to the best of our knowledge, no research has been carried out addressing dFC in cognitively unimpaired individuals with AD pathology markers.

Based on the presented premises and previous results from our group, indicating a sensitivity of the subfrequencies in the alpha band to early alterations, we aim to study how dFC values in the low (8-10 Hz) and high alpha ranges (10-12 Hz) change in relation to p-tau231, a non-invasive marker of AD and Aβ pathology. Furthermore, we aim to evaluate which brain areas are most affected by this relationship. We expect to find early dFC alterations associated with the individuals’ p-tau231 levels, even in the absence of AD symptomatology. Additionally, we will evaluate whether other risk factors contribute to the alteration observed in dFC.

## 2. Methods

### 2.1 Participants

The present study was carried out with a sample of 73 cognitively unimpaired individuals with available plasma p-tau231 determinations, extracted from a group of 140 participants with a valid MEG recording. Volunteers for this study are part of a larger initiative (“Study of the anatomo-functional connectome of AD-relatives: an early intervention on cognition and lifestyles”), aimed at longitudinally following the evolution of cognitively healthy at-risk and not-at-risk individuals over time in subsequent waves. The data used for the current study belongs to the second evaluation, during which the plasma p-tau231 determination was performed.

All participants were genotyped regarding their apolipoprotein E (APOE) carriage and had an available MRI scan acquired during the baseline assessment (2 to 3 years prior) that was used for this study. During this follow-up, all participants underwent an MEG scan, a blood sample extraction for plasma markers’ determination, and a thorough neuropsychological assessment to ensure a healthy cognitive status. Table 1 summarizes relevant demographic and clinical information of the sample.

**Table 1.**
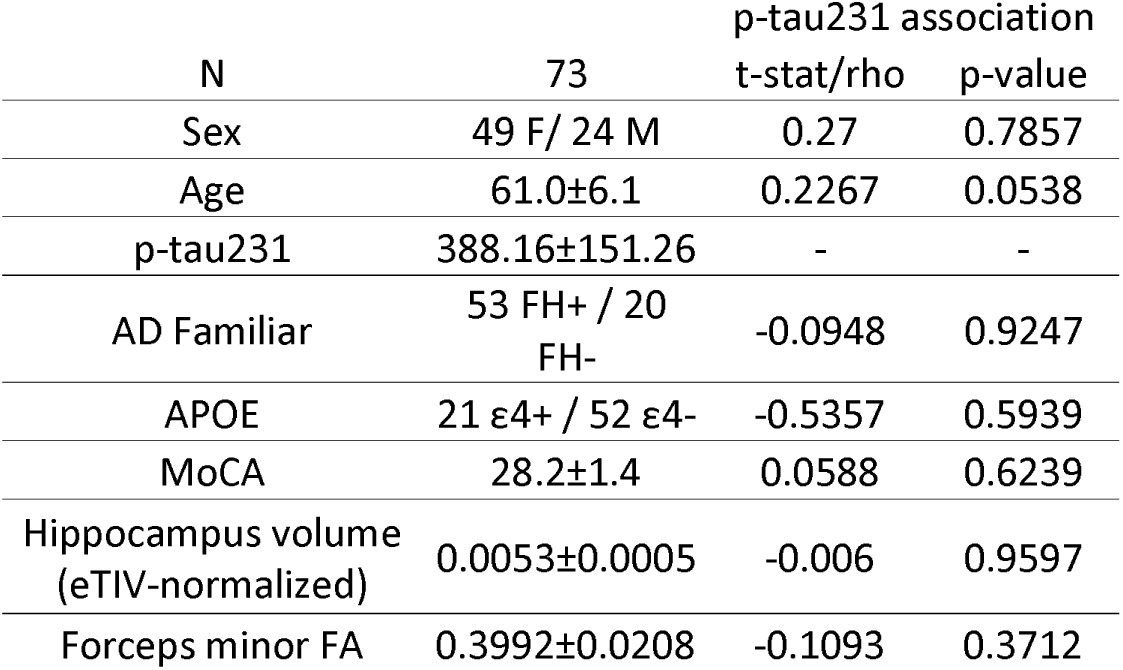
Sample demographics. The association with p-tau231 levels and the rest of the variables is shown in the two rightmost columns, displaying the independent samples t-test t-statistic and p-value for categorical values and Spearman correation’s rho and p-value for continuous variables. AD: Alzheimer’s disease; APOE: Apolipoprotein E; MoCA: Montreal cognitive assessment; FA: fractional anisotropy.

Exclusion criteria for the current study included: (1) history of psychiatric or neurological disorders or drug consumption in the last week that could affect MEG activity; (2) family history of dementia other than Alzheimer’s disease; (3) evidence of infection, infarction, or focal lesions in a T2-weighted MRI scan; (4) alcoholism or chronic use of anxiolytics, neuroleptics, narcotics, anticonvulsants, or sedative-hypnotics; (5) being younger than 50 or older than 80 years old; (6) a score in the Montreal cognitive assessment (MoCA) below 26; (7) unusable MEG recording or with less than 50 clean segments; (8) unavailable or unusable T1-weighted image; or (9) unavailable plasma p-tau231 determination.

The “Hospital Clínico San Carlos” Ethics Committee approved this study, and the procedure was performed following internationally accepted guidelines and regulations.

### 2.2 Plasma p-tau231 determination

Plasma p-tau231 concentration was quantified by competitive enzyme-linked immunosorbent assay, using a commercial kit (Human Phosphorylated Tau 231) (MyBioSource, Inc., USA, MBS724296), according to the manufacturer’s instructions. Tests were performed in duplicate, and an automated microplate reader (Biochrom Asys UVM340, Cambridge, UK) measured the optical density at 450 nm with MikroWin 2000 software (Berthold Technologies, Germany). Units are reported in pg/ml.

### 2.3 AD Family history

Family history of AD, which constitutes a compound risk for the disease [31], was defined as having a first degree relative (direct descendants or siblings) with confirmed Alzheimer’s disease. Relatives of patients were required to provide a medical report indicating the diagnosis of the patient following the NINCDS-ADRDA criteria [32].

### 2.4 Apolipoprotein E genetic evaluation

DNA was extracted from whole-blood samples. APOE haplotype was determined by analyzing single nucleotide polymorphisms (SNPs) rs7412 and rs429358 genotypes with TaqMan assays using an Applied Biosystems 7500 Fast Real-Time PCR machine (Applied Biosystems, Foster City, CA). A genotyping call rate over 90% per plate, sample controls for each genotype, and negative sample controls were included in each assay. Three well-differentiated genotyping clusters for each SNP were required to validate results. Intra- and interplate duplicates of several DNA samples were included.

### 2.5 Magnetic resonance imaging

Each subject’s T1-weighted MRI scan was acquired in a General Electric 1.5 T system. A high-resolution antenna was employed together with a homogenization Phased array Uniformity Enhancement filter (Fast Spoiled Gradient Echo sequence, TR/TE/TI = 11.2/4.2/450 ms; flip angle 12°; 1 mm slice thickness, 256 × 256 matrix and FOV 25 cm. The T1 images were processed using the FreeSurfer software (version 6.1.0) for automated cortical and subcortical segmentation and parcellation [33].

Additionally, diffusion weighted imaging (DWI) scans were acquired using a single shot echo planar imaging sequence with a TE of 96.1 ms, a TR of 12000 ms; a NEX 3 for increasing the signal-to-noise ratio, a FOV of 30.7 cm (plane resolution of 128 x 128) and a 2.4 mm slice thickness, yielding an isotropic voxel of 2.4 mm3. In the DWI sequence, one image had no diffusion sensitization (i.e., T2-weighted b0 images); and 25 were DWI (b = 900 s/mm2). DWI images were processed with AutoPtx for probabilistic tractography, as outlined in de Groot et al. [34]. The procedure extracted the mean fractional anisotropy (FA) for each tract.

The volumetric and tractography measures that were included in further analyses were parameters classically studied in the literature in the progress of AD: the total volume (mm3) of the bilateral hippocampi, normalized by the estimated intracranial volume (eTIV), and the FA of the forceps minor, which was chosen due to its relevance in the data-driven results obtained from the initial dFC analyses in this study.

### 2.6 MEG Data acquisition

Each participant underwent a five-minute, eyes-closed, resting-state MEG scan at the Center for Biomedical Technology in Madrid (Spain) using a 306-sensor Elekta Vectorview system (Elekta AB, Stockholm, Sweden). Ocular and cardiac activity was monitored with two sets of bipolar electrodes. Head position was continuously tracked with four head position indication coils placed on the participants’ forehead and mastoids. These coils, along with approximately 200 points of the head’s shape, were digitized using the PolhemusFASTRAK system (Polhemus, Colchester, VT, USA). The procedure was conducted in a magnetically shielded room, and participants were advised to remain as motionless as possible. The data were filtered online between 0.1 and 330 Hz and digitized at a rate of 1000 Hz. For data processing, the spatiotemporal signal space separation method [35], as implemented in the Neuromag software (MaxFilter version 2.2, with a correlation threshold of 0.90 and a correlation window of 10 seconds), was employed to eliminate external noise and adjust for any head movements during the MEG scan.

### 2.7 MEG preprocessing and source reconstruction

MEG data were blindly preprocessed by an electrophysiology expert using FieldTrip software [36]. This process involved dividing the continuous data into non-overlapping, artifact-free 4-second segments, with padding of 2 seconds of real data to each side of each segment. Participants who had at least 50 valid segments were selected for further analyses, given the high number of samples needed for entropy calculation (see next section). In addition, to improve the comparability across subjects, the number of segments was set to exactly 50, discarding data from those participants with more valid segments. Due to the high redundancy in the data after applying the signal space separation method [37], only data from magnetometers were used in further analyses.

The individual T1-weighted MRI scan was used for personalized source reconstruction. The source model was established in MNI space using a uniform 3D grid with 10 mm spacing, and source positions were identified based on the automated anatomical labeling atlas [38], resulting in 1210 cortical sources distributed across 80 regions of interest. This source model was then linearly transformed to each individual T1-weighted scan. The scans were segmented using SPM12 software [39], creating a brain mask, which was used to construct the participants’ single-shell realistic head models. This head model was combined with the individual source model for the generation of a lead field using a modified spherical solution [40].

To derive the time series for each source location, a linearly constrained, minimum variance beamformer was used as the inverse method. The sensor-space data was first filtered in the low alpha (8-10 Hz) and high alpha (10-12 Hz) frequency bands through a finite impulse response filter with an order of 1800, designed with a Hamming window. After filtering, the 2-seconds of padding were discarded, to avoid edge effects in the data. The beamformers were computed using the covariance matrix of the data filtered between 2 and 45 Hz, with a regularization level set at 1% of the average sensor power.

### 2.8 Functional connectivity

FC was estimated using the amplitude envelope correlation with leakage correction (AEC-c; [41]) between the time series of each pair of the 1210 cortical sources. After orthogonalization, the absolute value of the Pearson correlation between the time series’ envelopes was used. AEC was chosen because of its reliability, within and between subject consistency [42,43] and due to its sensibility to dFC in previous studies with similar objectives [22,30,44].

#### Static and dynamic functional connectivity calculation

We calculated AEC-c in each of the defined signal segments, and then averaged over those, obtaining the so-called static FC (Figure 1A), resulting in connectivity matrices of dimension 1210 sources × 1210 sources. Finally, we averaged the FC matrix across rows to obtain the connectivity nodal strength, which represents the average connectivity of each cortical source position with all the other sources (strAEC).

**Figure 1.**
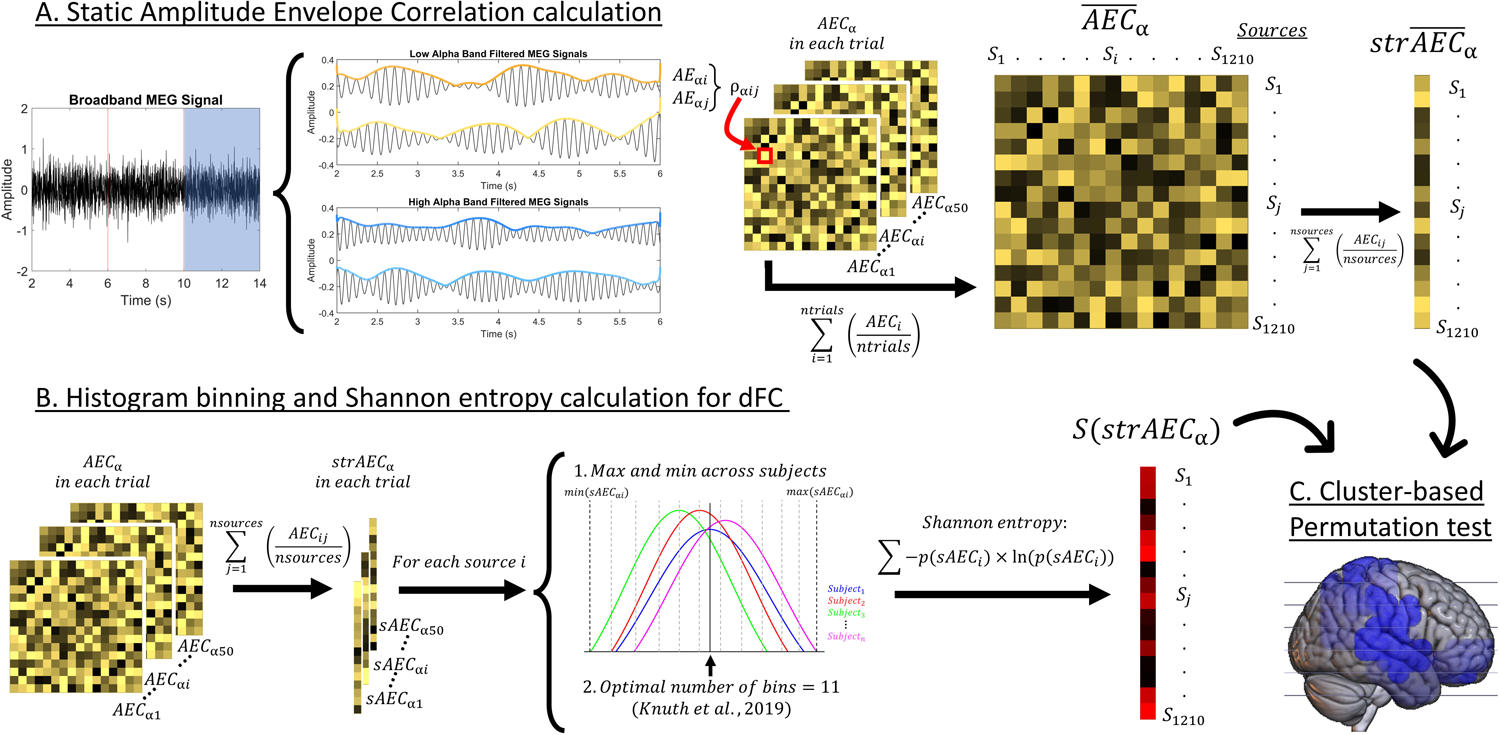
Depiction of the methods used in this study. (A) Description of the procedure to calculate the static AEC-c, including the signal filtering, envelope extraction through the Hilbert transform, correlation between envelopes, and averaging over trials and sources to obtain a measure of static strength (strAEC) for each source. (B) Description of the procedure to calculate the entropy for the nodal strength of each source position in each epoch. First, the nodal strength for each source position and epoch was calculated. We used a histogram with 11 bins to calculate Shannon entropy. This resulted in a value of entropy for each source position and participant (S(strAECα)), which accounts for their dFC. (C) A cluster-based permutation test is used to evaluate the correlation between the p-tau231 levels and the obtained dFC and sFC values.

For the sake of studying dFC, we calculated the entropy of the series of connectivity nodal strengths for each source position prior to the segment-averaging (Figure 1B). First, starting from the AEC-c connectivity matrices that were not averaged across time (dimensions 1210 sources × 1210 sources × 50 segments), we averaged these matrices over the second dimension, obtaining a temporal series of connectivity strengths for each cortical source position of dimensions 1210 sources × 50 segments. Afterwards, the entropy of each of those time series was calculated by means of histograms with linearly spaced bins.

Firstly, to determine the optimal number of bins for the creation of histograms in our dataset, we followed the guidelines exposed by Knuth [45] for each of the sources’ connectivity strength time series in all our participants. We kept the median of the resulting calculations, obtaining an optimal number of 11 bins. After that, and again to assure comparability between subjects, we defined the minimum and maximum values for the binning procedure in each cortical source by calculating the minimum and maximum value of the connectivity strength of a cortical source across all participants. Finally, following the optimal binning procedure, the Shannon entropy was calculated [46]:

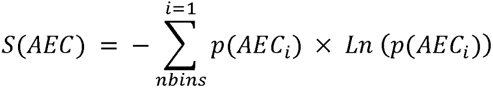

Where S is the entropy operator, AEC is our time series of connectivity strengths, *p(AEC_i_* is the probability for connectivity strength to fall under bin *i*, *Ln* is the natural logarithm and *nbins* is the total number of bins. The resulting entropy vectors included one value of entropy for each of the 1210 sources, representing the level of dynamicity of the cortical source connectivity over time. In other words, each source was assigned a value which portrays how variant the FC was over time, with higher values indicating a higher variability and vice versa. Figure 1 depicts the calculation process of FC parameters in this study.

### 2.9 Statistical analyses

Initially, demographic characteristics were evaluated using an independent samples t-test to compare p-tau231 levels between groups for categorical variables such as sex, APOE ε4 status, and AD family history. Spearman correlations were employed to assess the relationship between p-tau231 and quantitative variables, such as age, MoCA, eTIV-normalized hippocampal volume and FA at the forceps minor.

#### Spatial FC analysis

Statistical analyses were carried out in a data-driven, stepwise manner. Firstly, we used a cluster-based permutation test (CBPT; [47]) using a Montecarlo approach to assess which sources’ sFC and dFC strength in the high alpha and low alpha range were associated with p-tau231 levels. This nonparametric analysis accounts for multiple comparisons. Two-sided, partial correlations were used to assess the relationship between sFC or dFC and p-tau231, with sex and age as covariates. The source-level and cluster-level significance thresholds were set to α = 0.05. Significant sources were grouped based on their spatial contiguity, obtaining a cluster-level statistic defined as the sum of the individual statistics of each source within the cluster. We conducted 10000 permutations on the original data to create a surrogate distribution of random cluster-level statistics, which was then used to compare our original cluster size to the surrogate size distribution.

The clusters identified in the first step indicate brain areas showing significant correlation between sFC or dFC strength in general and levels of p-tau231. However, from this analysis alone, it is not possible to determine whether these clusters have different spatial patterns of connectivity. Therefore, we then performed a seed-based connectivity analysis, using the cluster identified in the first step as the seed. This results in a set of seed-based connectivity values for each of the source positions outside the seed. Again, we used a CBPT to identify significant partial correlations between sFC or dFC and p-tau231 levels, using sex and age as covariates. The result is a set of significant clusters of brain areas showing significant correlations with p-tau231 in their level of FC with the seed region. To obtain a single FC value for each subject, we calculated the average connectivity between the original cluster (seed) and the sources from the emerging cluster (secondary cluster), which we will refer to as the “seed-secondary cluster FC” from now on. Post-hoc Spearman partial correlation analyses were performed between the seed-secondary cluster dFC or sFC values and p-tau231 levels, using sex and age as covariates, to investigate the magnitude and directionality of the previous results.

#### Risk factor linear regressions

Finally, we investigated whether the observed changes in sFC or dFC were related to other risk factors aside from p-tau231 levels. To do so, we built three backwards stepwise linear regression analyses to explain the sFC or dFC of the previously found significant clusters:

● Firstly, we created a regression model including age, sex, presence of the APOE allele ε4, AD family history and MoCA scores. We also included sFC for dFC significant clusters and vice versa.
● Secondly, we added p-tau231 levels to the previous model to find out which parameters remained significant.
● Finally, an additional regression model was constructed from the previous one by adding factors classically studied in the continuum of AD: the eTIV-normalized hippocampal volume and fractal anisotropy of tracts related to the regions included in the clusters (i.e. the forceps minor) to the surviving factors of the previous stepwise regression analysis.

It is important to remark that we used backwards stepwise models as it was recommended in previous literature [48,49], given that the backward procedure the model is initiated with the full variable set, inspecting the significance of all possible variables, and that this procedure is better able to handle multicollinearity than a forward method.

## 3. Results

### 3.1 Sample demographics and variables of interest

Sample demographics and variables of interest used in this study are shown in Table 1.

The only variable that showed an almost significant correlation with p-tau231 was age (R = 0.2267, p-value = 0.0538), supporting the inclusion of this variable as covariate in the CBPTs of the study.

### 3.2 dFC increases along with p-tau231 values over orbitofrontal and temporal areas

While sFC strength values did not correlate significantly with p-tau231 values in any of the studied frequency bands, a cluster of significant correlation between dFC and p-tau231 emerged in the low alpha frequency band (p-value = 0.0460). This cluster showed a positive correlation between dFC nodal strength and p-tau231 levels, and included parts of the right temporal pole and right inferior, middle, and superior temporal gyri, along with the right parahippocampus (Figure 2, yellow).

**Figure 2.**
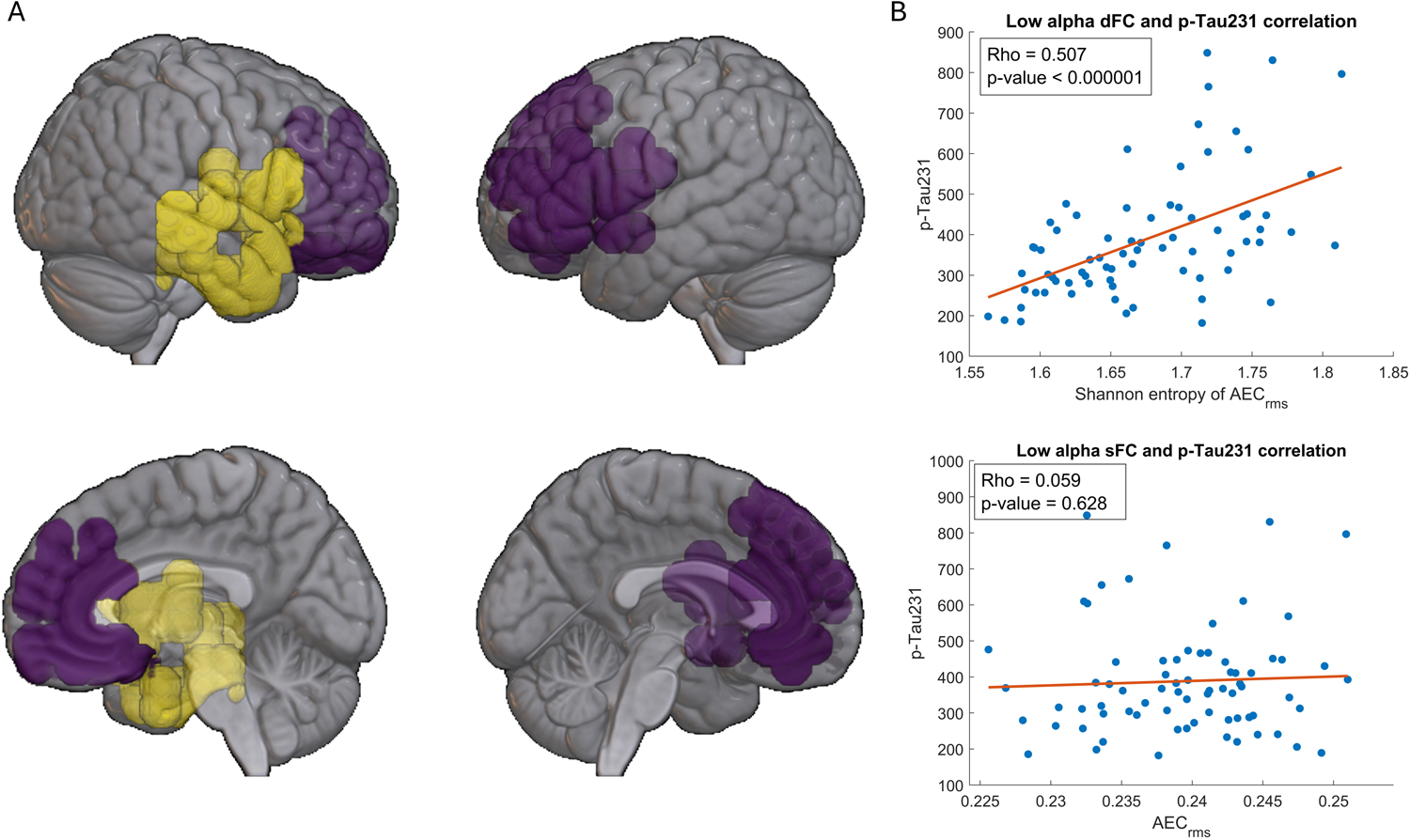
Clusters of significant association between dFC and p-tau231. A) The regions that show a positive correlation between their dFC levels and p-tau231 are represented in yellow (original cluster). Marked in purple are the regions that emerge after the second cluster-based permutation test (secondary cluster), whose dFC values with the original cluster show a positive correlation with p-tau231. B) Scatterplots and correlation values for the seed-secondary cluster dynamic functional connectivity (dFC; top) and static functional connectivity (sFC; bottom) with p-tau231 levels.

To better characterize this relationship, we studied the seed-based dFC of these sources with the rest of the brain in the same frequency band, using a single-sided contrast for a positive correlation between dFC and p-tau231. This analysis unveiled a unique cluster with a significant positive correlation between p-tau231 and the dFC from the original cluster (p-value = 0.0044). This group of sources included the right and parts of the left orbitofrontal cortices, left inferior and middle frontal gyri, bilateral anterior cingulate gyrus, and left parahippocampus (Figure 2, purple).

Spearman partial correlations between p-tau231 levels and the seed-secondary cluster dFC and sFC and p-tau231 revealed a strongly positive correlation between dFC and p-tau231 (Rho = 0.507, p-value = 6.3·10⁻⁶; Figure 2B), while the sFC did not significantly correlate with p-tau231 (Rho = 0.058, p-value = 0.6279; Figure 2C).

### 3.3 dFC is related to p-tau231 levels, sFC, and fractional anisotropy

Backwards stepwise linear regression models were built to explain the seed-secondary cluster dFC in the low alpha frequency band. Firstly, we evaluated the influence of the following variables on the dFC between the original and seed-based clusters: age, sex, MoCA scores, presence of the APOE allele ε4, AD family history, and sFC between clusters. The resulting model was significant (R-squared = 0.107, p-value = 0.0187), and only preserved two variables: age (p-value = 0.0735, β-value = 0.0021) and sFC (p-value = 0.0229, β-value = 2.77).

Subsequently, we built a second stepwise model by adding the levels of p-tau231 to find out which of the previously selected variables remained. The resulting model was significant (R-squared = 0.328, p-value = 9.0·10⁻⁷). This model discarded age as a relevant variable for the model, preserving p-tau231 (p-value = 1.6·10⁻⁶, β-value = 0.00021) and sFC (p-value = 0.0212, β-value = 2.43).

Finally, we added two additional factors to the previous one: eTIV-normalized hippocampal volume and FA at the forceps minor, given the location of the significant cluster. The resulting model was significant (R-squared = 0.389, p-value = 4.6·10⁻⁷) and preserved 3 variables: p-tau231 (p-value = 3.2·10⁻⁶, β-value = 0.00021), sFC (p-value = 0.0134, β-value = 2.65), and FA at the forceps minor (p-value = 0.0541, β-value = −0.569). A complete statistical report on the resulting linear regression models can be found in Supplementary Table S1.

The correlation of the significant variables and dFC can be found in Figure 2B for p-tau231 (R = 0.507, p-value < 0.0001), and in Figure 3 for sFC (R = 0.256, p-value = 0.0290) and the FA at the forceps minor (R = −0.270, p-value = 0.0250).

**Figure 3.**
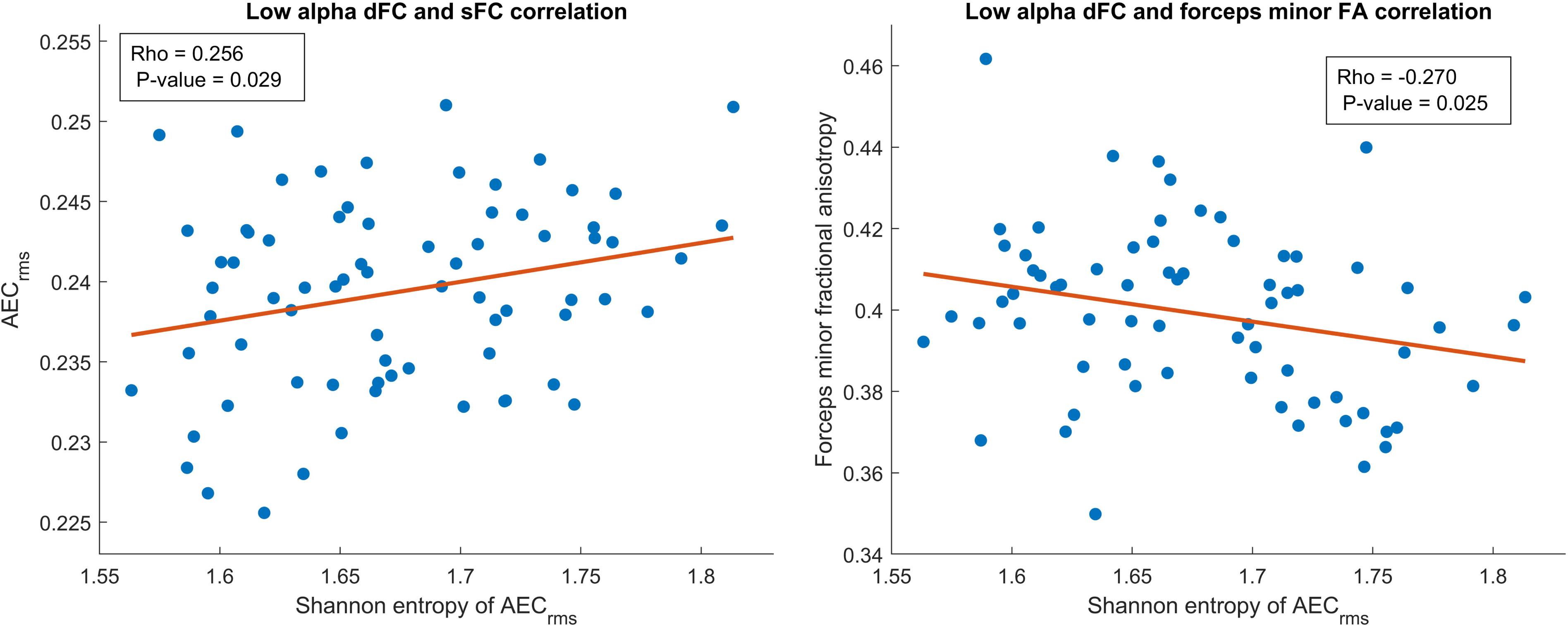
Correlation between dFC and linear regression model predictors. Scatterplot showing the positive association between dynamic functional connectivity (dFC) and static functional connectivity (sFC) of the seed-secondary cluster depicted in Figure 2 (left) and between dynamic functional connectivity of the seed-secondary cluster and the FA of the forceps minor (right).

## 4. Discussion

This study developed an unprecedented evaluation of the relationship between p-tau231 levels and both sFC and dFC through MEG. To this end, we calculated sFC through AEC-c and derived dFC from the entropy of the FC series over a sliding window, and calculated its spatial correlation with p-tau231 levels using non-parametric CBPTs. We found a cluster where dFC significantly and positively correlated with p-tau231 levels, located over the anterior parts of the temporal lobe and parahippocampal cortex, the orbitofrontal cortices, and bilateral anterior cingulate cortex. Moreover, we investigated which risk factors could be related to the changes observed in dFC through stepwise linear regression models. The linear regression models identified other factors influencing dFC, such as sFC and the FA of the forceps minor. These results complement and expand upon previous literature on the alterations of dFC in healthy individuals at risk of AD, addressing, for the first time, how this parameter changes in electrophysiological recordings.

The main finding from the present study is a cluster of positive correlation where the dFC values correlate with p-tau231 in the low-alpha band, as hypothesized based on previous literature [26,27]. No such results emerge when addressing sFC and p-tau231. This finding is of special relevance, as dFC could constitute a more sensitive metric than sFC to early alterations associated with AD pathology, due to the detection of subtle dynamic changes traditionally considered “noise” or interpersonal variability [50]. While previous fMRI studies in healthy individuals with AD pathology yield inconsistent results regarding the directionality of dFC changes, studies carried out addressing the prodromal and dementia stages consistently point towards a reduction of dFC, both with MEG and fMRI [23,24,51]. This decline is hypothesized to be another sign of the disconnection and the excitation/inhibition imbalance that characterize AD and underlie a loss of coupling strength that leads to a decrease of dFC [22]. While our results could seem inconsistent with such a trend, sFC is also known for not following a linear decline, but rather an inverted U-shaped pattern, marked by a state of hyperconnectivity in the early stages of Aβ accumulation, and hypoconnectivity later on in the continuum. The initial hyperconnectivity is caused by an underlying state of neuronal hyperexcitability that contributes to increased Aβ deposition which, in turn, increases hyperexcitability [52,53]. In this line, previous studies by our group carried out with p-tau231 find a positive association with sFC along with centrality increases in posterior regions as predicted by theoretical and computational models of the early stages of AD [54–56]. Therefore, dFC could follow an equivalent trend and experience an increase in the early stages of the disease in association with Aβ increases. Nevertheless, future research should be done to specifically address the underlying cause of the dFC increases.

The areas forming the original cluster include areas such the right temporal pole and right inferior, middle, and superior temporal gyri, along with the right parahippocampus, which show a positive relationship between dFC and p-tau231. The secondary cluster that emerged from the seed-based analysis of the original one includes regions such as the right and parts of the left orbitofrontal cortices, left inferior and middle frontal gyri, bilateral anterior cingulate gyrus, and left parahippocampus, whose dFC with the original cluster is positively associated with p-tau231. These results are congruent with previous literature, as they suggest early dFC alterations that span regions from the DMN, which is known to be especially vulnerable to AD pathology, as well as the orbitofrontal cortex, which accumulates Aβ from the earliest stages of the continuum [23,28,57]. Moreover, a previous study carried out by Lorenzini et al. [58] involves similar areas. In this study of 736 healthy individuals, they evaluated the relationship between CSF Aβ levels and centrality variability. They report a decrease of centrality variability in a frontotemporal cluster encompassing areas similar to the ones that emerge in the present study. While the spatial distribution of both sets of results seems to suggest early alterations in the dynamic of frontotemporal regions, the comparability of the meaning of both sets of results in combination is hampered. More research is needed to understand the underlying neuronal cause of dFC increases and graph-theory dynamics, along with studies that link electrophysiological and fMRI dFC changes. While electrophysiological techniques might be more suitable to study dFC, fMRI has been more widely used, but results are difficult to interpret due to physiological artifacts, variation over time in both the mean and variance of the blood-oxygen-level dependent signal [59], and different methodological differences such as the technique used to estimate dFC or the time-scale between studies. Previous investigations on multiscale entropy to assess signal complexity in AD showed that the directionality of the results depended on the time-scale for the parameter calculation, finding increased signal complexity in controls for short time-scales, and decreased for larger time-scales [60,61]. Consequently, comparing results from different studies is intricate, and multi-analysis studies are needed to better understand the relationship between different dFC metrics and improve comparability [62].

To better understand which factors other than p-tau231 levels influence dFC, we performed stepwise regression models. The initial model identified age and sFC as significant predictors of dFC levels. However, incorporating p-tau231 into the model resulted in the exclusion of age and enhanced the explanatory power of the model, with R-squared values increasing from 0.107 to 0.328. These findings suggest that although related, p-tau231 has an additional pathological component to that of aging, thus having a greater impact over dFC. Subsequently, two structural and classically used pathology parameters in the context of AD were included in the model: normalized hippocampal volume and FA. Specifically, we added the FA of the forceps minor due to the location of the areas identified through the CBPTs. This resulted in the maintenance of sFC and FA of the forceps minor as predictors, alongside p-tau231, increasing the resulting R-squared of the model to 0.389. Notably, their influences on dFC are independent of p-tau231 and show significant associations with dFC both when included in the model and when addressed through separated correlation analyses: a direct association for sFC and an inverse one for the FA of the forceps minor (Figure 3). The inclusion of these plasma, functional, and structural variables in the models, as well as the direction of their association with dFC, suggest that the increase observed in relation to p-tau231 in frontotemporal regions is of pathological nature and begins early on. Prior research indicates that sFC increases during the initial phases of AD and in association with p-tau231 [10], and extensive literature links FA reductions with AD progression [63]. Our findings highlight a complex interplay among these biomarkers, with dFC being the only parameter correlating significantly with p-tau231. This underscores the importance of studying brain dynamics as a potential early biomarker for detection of individuals entering the AD continuum.

Some limitations must be noted. While this study seeks to develop the knowledge around the initial stages of AD, and our study is a pioneer in the follow-up of high-risk individuals within the specific age range where early pathological signs are expected to occur, a longer follow-up period is required to correctly classify cases and controls. Relevantly, this study is part of an initiative aimed at tracking these individuals for decades, with the third time point assessment already approved and funded. Further, future assessment rounds should include other standardized and widely accepted techniques to directly quantify AD pathology to adequately evaluate the relationship between connectivity parameters and the levels of pathology markers. Nonetheless, plasma p-tau231 might represent an alternative and more ecological marker that is also less invasive, widely accessible to researchers and clinicians, and more cost-efficient. On the other hand, the use of a sliding-window of a fixed size does not come without disadvantages, given that recent approaches have demonstrated that dFC can be well-described at a variety of time-scales, and there may be a mismatch between the temporal scale of the underlying fluctuations and the predefined fixed window length. However, previous literature addressing the frequency bands under study in the present work have described relevant sFC features in the AD continuum [10,11,14] with 4-second windows, which have also been recommended for the estimation of fluctuations in FC of resting state networks [43]. Finally, more research is needed, combining electrophysiological measures and fMRI, as well as using different dFC estimation techniques, in order to get a better understanding of the underlying neural alterations that induce the dFC changes.

Overall, our study takes a pioneering approach to study alterations in the dynamics of brain connectivity associated with the plasma pathology marker, p-tau231, in cognitively unimpaired individuals. Our results are innovative, as this is the first work to address dFC in healthy individuals using plasma markers and MEG. Moreover, they are consistent and expand upon previous literature, suggesting that changes associated to AD pathology can be observed very early on using dFC, which might be more sensitive to subtle alterations than sFC, where these might be lost in the averaging process. Lastly, this early increase in dFC in frontotemporal regions appears to have a pathological nature given its relationship with other plasma, functional and structural measures.

## 5. Data availability

The data and the algorithms that support the findings of this study are available from the corresponding author, upon reasonable request.

## 6. Acknowledgements

We would like to thank all the participants that have selflessly given us their time and made this study possible.

## Supplementary materials

**Table S1.**
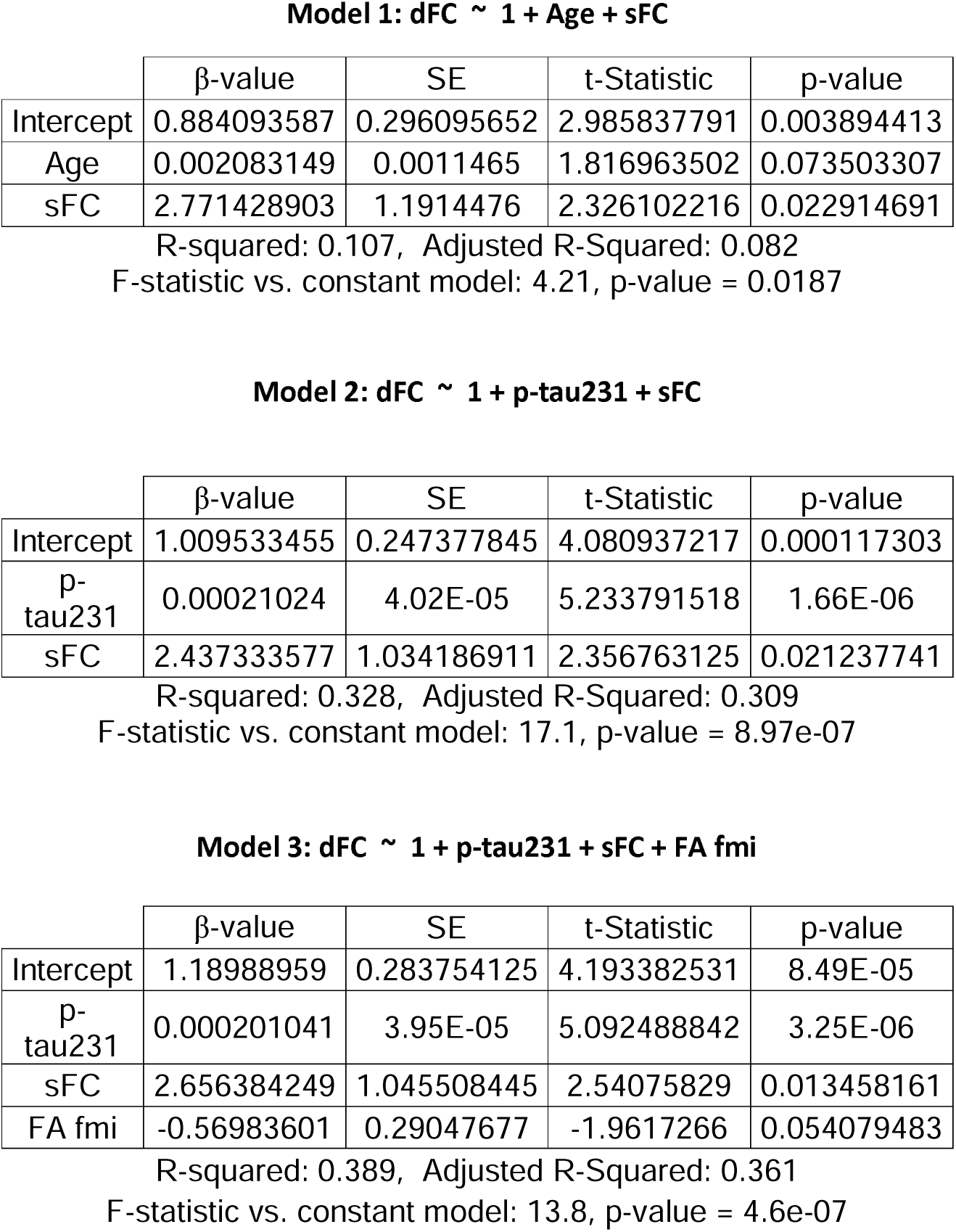
Linear regression results for the proposed models. SE stands for standard error, sFC for static functional connectivity and FA fmi for fractional anisotropy of the forceps minor.

