## Supplementary material for "Dynamic functional connectivity is modulated by the amount of p-Tau231 in blood in cognitively intact participants": Table 1

**Table 1. Sample demographics.** The association with p-tau231 levels and the rest of the variables is shown in the two rightmost columns, displaying the independent samples t-test t-statistic and p-value for categorical values and Spearman correation’s rho and p-value for continuous variables. AD: Alzheimer’s disease; APOE: Apolipoprotein E; MoCA: Montreal cognitive assessment; FA: fractional anisotropy.

|  |  | p-tau231 association | |
| --- | --- | --- | --- |
| N | 73 | t-stat/rho | p-value |
| Sex | 49 F/ 24 M | 0.27 | 0.7857 |
| Age | 61.0±6.1 | 0.2267 | 0.0538 |
| p-tau231 | 388.16±151.26 | - | - |
| AD Familiar | 53 FH+ / 20 FH- | -0.0948 | 0.9247 |
| APOE | 21 ε4+ / 52 ε4- | -0.5357 | 0.5939 |
| MoCA | 28.2±1.4 | 0.0588 | 0.6239 |
| Hippocampus volume (eTIV-normalized) | 0.0053±0.0005 | -0.006 | 0.9597 |
| Forceps minor FA | 0.3992±0.0208 | -0.1093 | 0.3712 |
