## Supplemental Table 1 for "Dynamic functional connectivity is modulated by the amount of p-Tau231 in blood in cognitively intact participants"

**Table S1**. Linear regression results for the proposed models. SE stands for standard error, sFC for static functional connectivity and FA fmi for fractional anisotropy of the forceps minor.

**Model 1: dFC ~ 1 + Age + sFC**

|  | β-value | SE | t-Statistic | p-value |
| --- | --- | --- | --- | --- |
| Intercept | 0.884093587 | 0.296095652 | 2.985837791 | 0.003894413 |
| Age | 0.002083149 | 0.0011465 | 1.816963502 | 0.073503307 |
| sFC | 2.771428903 | 1.1914476 | 2.326102216 | 0.022914691 |
| R-squared: 0.107, Adjusted R-Squared: 0.082 F-statistic vs. constant model: 4.21, p-value = 0.0187 | | | | |
| **Model 2: dFC ~ 1 + p-tau231 + sFC** | | | | |
|  | β-value | SE | t-Statistic | p-value |
| Intercept | 1.009533455 | 0.247377845 | 4.080937217 | 0.000117303 |
| p-tau231 | 0.00021024 | 4.02E-05 | 5.233791518 | 1.66E-06 |
| sFC | 2.437333577 | 1.034186911 | 2.356763125 | 0.021237741 |
| R-squared: 0.328, Adjusted R-Squared: 0.309 F-statistic vs. constant model: 17.1, p-value = 8.97e-07 | | | | |
| **Model 3: dFC ~ 1 + p-tau231 + sFC + FA fmi** | | | | |
|  | β-value | SE | t-Statistic | p-value |
| Intercept | 1.18988959 | 0.283754125 | 4.193382531 | 8.49E-05 |
| p-tau231 | 0.000201041 | 3.95E-05 | 5.092488842 | 3.25E-06 |
| sFC | 2.656384249 | 1.045508445 | 2.54075829 | 0.013458161 |
| FA fmi | -0.56983601 | 0.29047677 | -1.9617266 | 0.054079483 |
| R-squared: 0.389, Adjusted R-Squared: 0.361 | | | | |
| F-statistic vs. constant model: 13.8, p-value = 4.6e-07 | | | | |
